# Visceral mesoderm signaling regulates assembly position and function of the *Drosophila* testis niche

**DOI:** 10.1101/2021.09.08.459436

**Authors:** Lauren Anllo, Stephen DiNardo

**Affiliations:** Perelman School of Medicine at the University of Pennsylvania, The Penn Institute for Regenerative Medicine

**Keywords:** niche, stem cell, *Drosophila*, testis, visceral mesoderm, *biniou*, Slit, FGF, *islet*

## Abstract

Tissue homeostasis often requires a properly placed niche to support stem cells. Morphogenetic processes that position a niche are just being described. For the *Drosophila* testis we recently showed that pro niche cells, specified at disparate positions during early gonadogenesis, must assemble into one collective at the anterior of the gonad. We now find that Slit and FGF signals emanating from adjacent visceral mesoderm regulate assembly. In response to signaling, niche cells express *islet*, which we find is also required for niche assembly. Without signaling, niche cells specified furthest from the anterior are unable to migrate, remaining dispersed. Function of such niches is severely disrupted, with niche cells evading cell cycle quiescence, compromised in their ability to signal the incipient stem cell pool, and failing to orient stem cell divisions properly. Our work identifies both extrinsic signaling and intrinsic responses required for proper assembly and placement of the testis niche.

## Introduction

Stem cells play a vital role in tissue repair and maintenance, and their loss is associated with degeneration. To maintain stem cells within any tissue, these cells must receive self-renewal signals, often from their resident niche, a microenvironment that supports and directs stem cell behavior (Losick et al., 2011; Moore and Lemischka, 2006; Morrison and Spradling, 2008). Assembly of a niche is crucial for stem cell function, and positioning of the assembled niche in the appropriate location during organ development ensures that niche signals remain accessible and confined to stem cells. Regulation of niche assembly and position are therefore relevant to tissue homeostasis. Morphogenetic and signaling events that underly formation of tissues where stem cells reside are being described in tissues such as the intestinal crypt and hair follicle (Greicius and Virshup, 2019; Kaestner, 2019; Martino et al., 2021; Rompalos and Greco, 2014; Shwartz et al., 2020). These tissues and others exhibit a paradigmatic compartmentalization of niche cells during organogenesis. Yet, how niche cells assemble in the appropriate position within their resident tissue remains largely unknown.

We study the *Drosophila* testis niche, which is well defined and has served as a paradigm for understanding niche-stem cell interactions. We refer to the testis niche, or niche cells, as the cells that emit signals to support neighboring stem cells. Our recent work pioneered live-imaging formation of this niche, enhancing its strength as a model (Anllo et al., 2019; Nelson et al., 2020). Appropriate placement of this niche is important for polarizing the testis, and enabling tissue function (Fuller, 1993; Lee et al., 2008; Tanentzapf et al., 2007). The niche resides at the apex of a closed tube, and directs germline stem cell (GSC) divisions such that some daughter cells are displaced from self-renewal signals. This arrangement facilitates the movement of differentiating cells further along the tube to eventually release mature sperm at the base (Fuller, 1993; Hardy et al., 1979; Kiger et al., 2001; Tulina and Matunis, 2001; Yamashita et al., 2003; Yamashita et al., 2007). Anchorage of the niche at the testis apex ensures proper niche positioning throughout the life of the fly. Without anchoring, the niche drifts from the apex, fails to properly orient divisions, and is eventually lost (Lee et al., 2008; Papagiannouli et al., 2014; Tanentzapf et al., 2007). Additionally, flies with defects in niche anchoring have reduced fertility (Lee et al., 2008), confirming the importance of niche position in testis function. While we have some knowledge of how niche positioning is maintained, how the niche is initially assembled in its correct position is unknown and is the focus here (Anllo et al., 2019; Lee et al., 2008; Papagiannouli et al., 2014; Tanentzapf et al., 2007).

The niche assembles at the gonad anterior during embryonic development in the male (Aboim, 1945; Le Bras and Van Doren, 2006; Sheng et al., 2009; Sinden et al., 2012). The gonad is spherical, comprised of germ cells (GCs) intermingled with and encysted by somatic gonadal precursor cells (SGPs) (Aboim, 1945; Jenkins et al., 2003). Prior to niche formation, prospective (pro) niche cells are specified from a subset of SGPs by coordination of Notch and EGFR signaling (Kitadate and Kobayashi, 2010; Okegbe and DiNardo, 2011). Once specified, pro niche cells undergo two phases of niche morphogenesis, namely a loose assembly as a cap at the gonad anterior, followed by compaction into a tight, spherical structure (Anllo et al., 2019; Le Bras and Van Doren, 2006). The process by which pro niche cells assemble at the anterior is dynamic. Pro niche cells are initially intermingled with germ cells (GCs). They extend protrusions to pull themselves onto the gonad periphery, and then migrate anteriorly along extracellular matrix (ECM) until they associate in a cap (Anllo et al., 2019). This cap assembles at a pole directly opposite a group of male specific somatic cells, msSGPs, located at the gonad posterior (Anllo et al., 2019; DeFalco et al., 2003). Once assembled at the anterior, the niche displays distinguishing markers of adhesion and gene expression including Fasciclin III (Fas3), E-cadherin (Ecad), and *unpaired (upd)* (Le Bras and Van Doren, 2006). Niche morphogenesis is complete at the end of embryogenesis, Stage 17, and this arrangement is preserved through development, compartmentalizing the niche to the tip of the adult testis. (Anllo et al., 2019; Sheng and Matunis, 2011; Sinden et al., 2012; Tanentzapf et al., 2007).

Our previous imaging showed that as niche assembly occurs the position adopted is tilted towards interior regions of the embryo (Anllo et al., 2019). The tilt suggested that tissues external to the gonad might be signaling to direct niche placement. The midgut is one tissue located near the assembled niche (Anllo et al., 2019). The midgut is surrounded by musculature derived from visceral mesoderm (Vm), and prior work suggested that Vm is a signaling center directing morphogenesis of the endoderm, salivary glands, and longitudinal visceral muscles (Azpiazu and Frasch, 1993; Bradley et al., 2003; Cimbora and Sakonju, 1995; Immerglück et al., 1990; Kadam et al., 2012; Tepass and Hartenstein, 1994). We thus suspected that the Vm could direct anterior assembly of the gonad niche.

Most of the Vm derives from segmentally repeated groups of mesodermal cells along the anterior-posterior axis of the embryo, specified by transcription factors including *bagpipe* and *biniou (bin).* After specification Vm precursors contact one another in lateral arrangements on either side of the endoderm (Azpiazu and Frasch, 1993; Zaffran et al., 2001). Vm cells undergo fusion to form the circular muscles that later surround the gut and direct its morphogenesis (Immerglück et al., 1990; Klapper et al., 2002; San Martin et al., 2001; Tepass and Hartenstein, 1994). Longitudinal muscles overlay the circular muscles, and derive from caudal Vm (cvm) precursors specified at the embryo posterior that migrate anteriorly over the Vm (Zaffran et al., 2001). Collectively, Vm tissue is known to express numerous signals, including Slit and FGFs. Slit activates Robo receptors, which act in cell adhesion and axon guidance during development. The FGFs Pyramus (Pyr) and Thisbe (Ths) activate the FGF receptor Heartless (Htl), which is important for guiding migration of cvm precursors over trunk Vm (Stathopoulos et al., 2004).

We reveal that Vm signals Slit and FGF are required to assemble a compartmentalized niche in the gonad. These signals are required for niche cell cytoskeletal polarity and anterior movement of pro-niche cells. In response to these signals, niche cells express the transcription factor *islet* (or *tup),* which is important for expression of axon guidance receptors in the nervous system (Santiago and Bashaw, 2014; Santiago and Bashaw, 2017; Yang et al., 2009). We demonstrate that Islet is also required to assemble the niche. Finally, we show that anterior niche assembly is important for proper niche function and behavior. Taken together, this work unveils how niche position arises during development.

## Results

### Visceral mesoderm is required for niche assembly and positioning

To test for a role of the Vm in positioning the niche, we examined gonads dissected from *biniou* mutant embryos, which lack Vm tissue. *biniou* encodes a FoxF transcription factor essential for Vm development, with expression reported solely in Vm precursors (Azpiazu and Frasch, 1993; Zaffran et al., 2001). In sibling controls dissected at the end of embryogenesis, Stage 17, when niche morphogenesis is normally complete, we observed a single anterior niche using both Fas3, a cell adhesion marker for niche cell boundaries, and *upd*>GFP, a marker for niche cell-specific gene expression (**Fig1A,C**). Anterior niche position was confirmed relative to male specific somatic gonadal precursor cells (msSGPs) at the gonad posterior. In contrast, gonads from *biniou* mutants often exhibited dispersed aggregates of niche cells (**Fig1B,D**). To rule out changes in the number of niche cells specified, we quantified niche and other somatic gonadal precursor cells (SGPs), and observed no differences between *biniou* mutant gonads and sibling controls (**Fig1G**). Thus, the dispersed niche phenotype in *biniou* mutants results from defects in niche assembly. These data indicate that *biniou* is required for anterior niche assembly, which in turn suggests that the Vm is required to position the niche.

**Figure 1.**
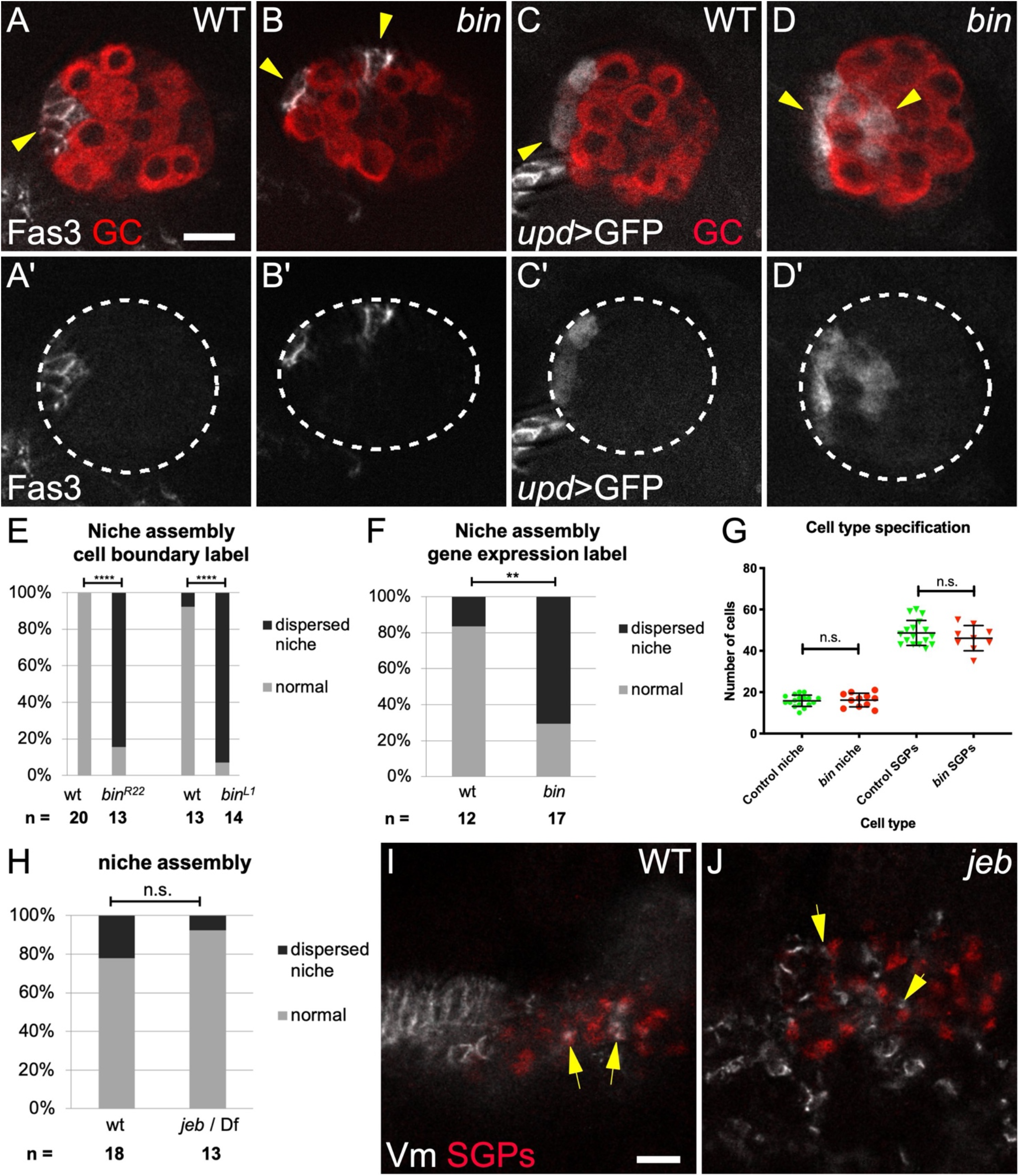
Visceral mesoderm is required for niche assembly and positioning. (**A**) Control stage 17 gonad when niche morphogenesis is complete, immunostained with Vasa (red, germ cells) and Fas3 (white, niche cells). (**B**) *biniou* mutant; Vasa (red) and Fas3 (white) reveal a dispersed niche (arrowheads). (**C**) Control and (**D**) *biniou* mutant gonads immunostained with Vasa (red) and expressing *upd*-Gal4, UAS-GFP in niche cells (white). (**A’-B’**) Fas3 alone; (**C’-D’**) GFP alone. Dotted lines, gonad boundary. (**E-F**) Quantification using (**E**) Fas3 or (**F**) *upd*>GFP as marker (p < 0.001, p = 0.004, respectively, Fisher’s exact test). (**G**) Number of niche and non-niche cells specified in *biniou* mutants compared to siblings. (**H**) Niche assembly is not affected in *jeb* mutants compared to siblings. (**I**) Control and (**J**) *jeb* mutant embryos (Stage 13, before gonad coalescence); arrows show SGPs (Traffic jam, red) in contact with Vm cells (Fas3, white). *jeb* mutants have a different arrangement of Vm precursors. Scalebars, 10 um.

### Visceral mesoderm tissue is required before niche assembly

To investigate the timing of the requirement, we examined *jelly belly* (*jeb*) mutants in which Vm cells are initially specified but do not complete development, and the Vm is missing by Stage 15 (Stute et al., 2004; Weiss et al., 2001). Stage 15 is the embryonic stage when niche assembly begins. Interestingly, *jeb* mutants had a normal number of niche cells and normal niche morphology (**Fig 1H**, and data not shown). This result, along with the lack of proper niche assembly in *biniou* mutants suggests that the Vm is necessary early, and dispensable by the time the niche assembles. Consistent with this, we also observed that at early stages prior to niche assembly, Vm precursor cells were intermingled with SGPs (**Fig 1I**). Such intermingling was also seen in *jeb* mutants (**Fig 1J**). In contrast, in *biniou* mutants we never observed intermingling of SGPs with the rare Vm precursors (**Fig S1**). These findings imply that Vm signals are active before commencement of niche assembly and may well involve direct cell contact between pro niche and Vm cells.

### Slit and the FGF ligands *pyr* and *ths* promote anterior niche assembly

*biniou* was reported to be expressed only in Vm, and only to affect its development (Zaffran et al., 2001). We were surprised to observe some Biniou protein accumulation in SGPs in coalesced gonads (**Fig S2**). We thus sought to confirm a role for the Vm in niche assembly by mining existing literature for Vm-expressed genes that encoded ligands. The two FGF ligands, *pyramus* (*pyr*) and *thisbe* (*ths*), which often act redundantly, met these criteria (Kadam et al., 2012; Stathopoulos et al., 2004). We confirmed that each was expressed in Vm cells (**Fig S3 A-B, D-E**). We observed expression in some other mesodermal cells outside the gonad, but not in the region where SGPs are located, interspersed among germline cells. In embryos where both *pyr* and *ths* were removed, gonadal niche cells were often dispersed, or assembled but not located at the gonad anterior (**Fig 2B, 2C, E**). A subset of Vm cells, the caudal visceral mesoderm (cvm), is missing in *pyr* and *ths* mutants, raising a possibility that these Vm ligands might act indirectly, through cvm, rather than directly in positioning the niche. However, niche placement is normal in a mutant lacking cvm (**Fig S4 E-F**). In fact, consistent with a direct ligand requirement in positioning the niche, we observed expression of the Heartless FGF Receptor in SGPs (**Fig S4 C-D**), and *Htl* mutants also exhibited niche assembly defects (**Fig 2F**). These results demonstrate that the FGF ligands *pyr* and *ths* are important for assembling an anterior niche and could emanate from the Vm to do so.

**Figure 2.**
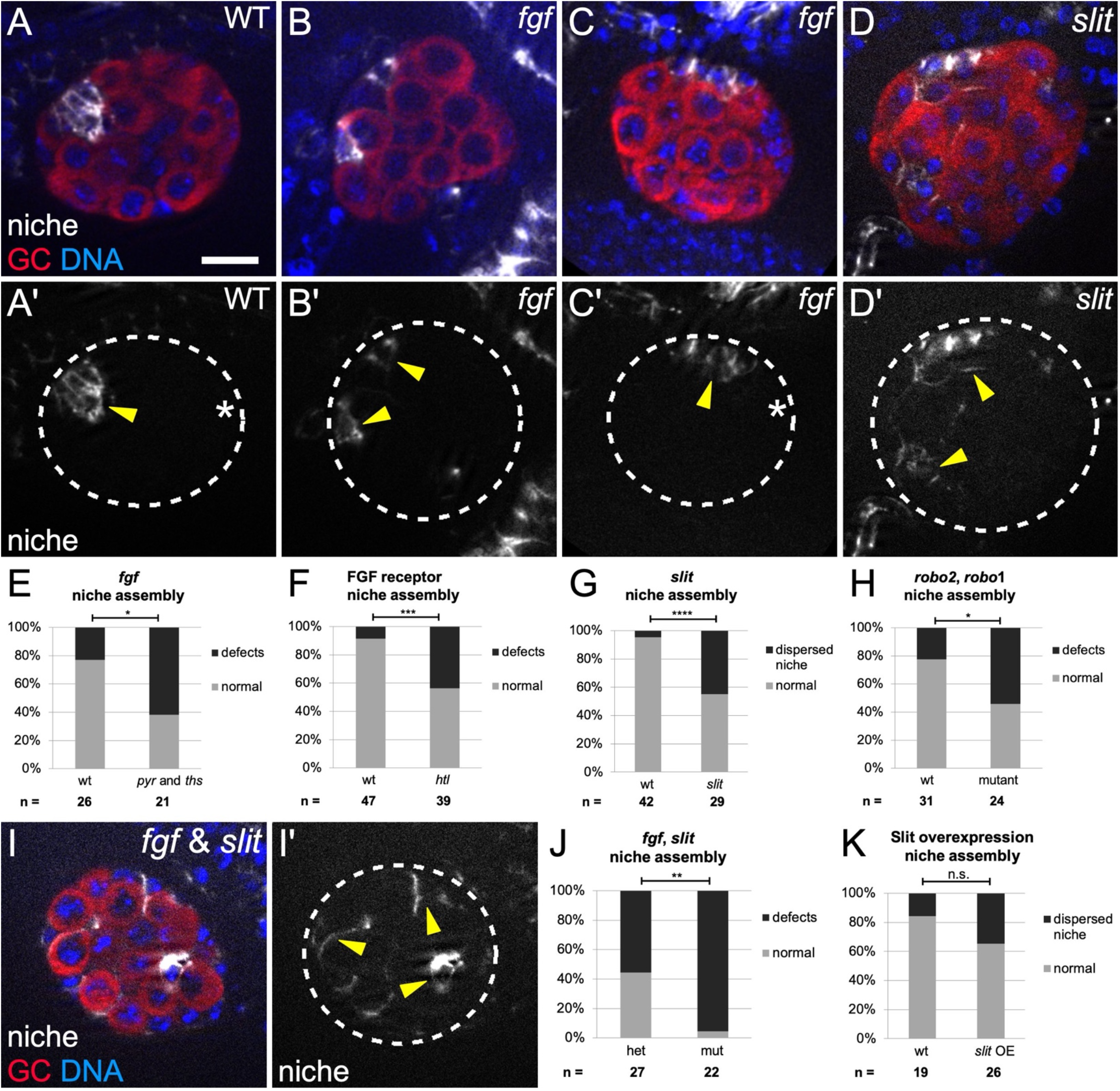
Slit and FGF signaling promote anterior niche assembly. (**A-D,I**) Stage 17 gonads, merge of Vasa (red, germ cells), Hoechst (blue, DNA), and Fas3 (white, niche cells), or single channel (Fas3); Dotted line, gonad boundary. Scale bar, 10 um. Prime panels show Fas3 (niche cells, arrowheads) alone. (**A**) Sibling controls have a single, anterior niche. (**B-C**) Df(2R)BSC25 gonads, with a deletion removing *pyr* and *ths* genes, exhibit niche defects such as (**B**) dispersed niche cell aggregates, and (**C**) niches not at the gonad anterior (asterisk, gonad posterior). (**D**) *slit[2]* mutant gonads often have dispersed niche cell aggregates. (**E-H**) Quantification of niche defects (Fisher’s exact test), (**E**) with *pyr* and *ths* removed (*fgf*) (p = 0.016), (**F**) FGF *htl* receptor mutant (p = 0.0003), (**G**) *slit* mutant (p < 0.0001), and (**H**) *robo2, robo1* double mutant (p = 0.024). (**I, J**) Combined mutant with *slit,* and *pyr* and *ths* removed (*fgf*) exhibits dispersed niche cells (p = 0.003). (**K**) Niche morphogenesis defects were not significant (n.s.) in Slit overexpression embryos.

The ligand Slit is expressed in Stage 13 Vm, and we confirmed this for Stages 13 and 16, before and during niche assembly, respectively (**Fig S3 C,F**) (Kraut and Zinn, 2004; Rothberg et al., 1990; Sandmann et al., 2006; Soplop et al., 2012). We also detected occasional expression in other mesodermal cells, some of which flank the forming gonad, but were not interspersed among germ cells, and thus not SGPs (**Fig S3 C, F**). We next examined *slit* mutants, which were previously shown to have a partially penetrant defect during an earlier phase of gonad development (Weyers et al., 2011). For that reason, we only analyzed that fraction of *slit* mutant gonads that formed properly, having bypassed the earlier role for Slit. In this manner we ensured that any effects on niche morphology were unlikely to be secondary to some block in proper gonad formation. Indeed, we observed niche assembly defects in up to 40% of such *slit* mutant gonads (**Fig 2D,G**). Consistent with a role for Slit, the Slit receptors Robo1 and Robo2 have been observed in SGPs (Weyers et al., 2011), and we detected niche morphogenesis defects in *robo1, robo2* double mutants (**Fig 2H; Fig S4 G-H**). These data support the idea that Slit, which is expressed in the Vm, contributes to proper assembly of an anterior gonad niche.

Since the removal of either Slit or the pair of FGF ligands resulted in a partial phenotype, we hypothesized that each of these pathways might independently contribute to niche assembly. Indeed, simultaneously removing Slit and both FGF ligands resulted in a virtually fully penetrant niche assembly defect (compare **Fig 2I, J** to **Fig 2E and G**). These data indicate that FGF and Slit act in parallel to facilitate niche assembly. Finally, we also observed defects in gonads from embryos heterozygous for *slit, pyr* and *ths* (**Fig S4 I-K**), suggesting that the dosage of signaling ligands is relevant for proper niche assembly. To summarize, mutants for *biniou,* which have no Vm, exhibit niche assembly defects, and the removal of two classes of signaling ligands, which each appear to emanate from the Vm, exhibit virtually identical niche assembly defects. We conclude that the Vm is the main tissue responsible for assembling the gonadal niche in its correct position.

Slit and FGF ligands usually direct migratory paths during morphogenesis. To test whether these ligands might be playing a directional role for niche assembly, we misexpressed each ligand broadly in mesoderm, and asked if that changed the position of niche assembly. Unfortunately, as seen before, *ths* or *pyr* overexpression led to general morphogenetic defects and impeded gonad formation such that no conclusion could be drawn (data not shown) (Sun and Stathopoulos, 2018). In contrast, *slit* overexpression occasionally yielded properly formed gonads. As with *slit* mutants, we only analyzed niche morphology where there was a properly coalesced gonad, to ensure that any effects on the niche were not secondary to some block to gonad formation. Surprisingly, niche morphogenesis was unaffected by Slit overexpression when compared to siblings (**Fig 2K**), suggesting that Slit is acting as a competence factor and not a directional cue for niche assembly. Interestingly, GFP-tagged Slit appeared to accumulate at the gonad periphery, likely in extracellular matrix (**Fig S3 G**). This unpolarized accumulation could be consistent with the idea that Slit is not a directional cue.

### Visceral mesoderm is required for anterior movement of pro niche cells

We showed previously that proper niche assembly involves several steps, the first of which necessitates that pro niche cells sort out of the internal milieu and onto the gonad periphery (Anllo et al., 2019). Signals from the the Vm are not required for this step as niche cells in both control and *biniou* mutants were located at the gonad periphery to a similar degree (**Fig 3G**).

**Figure 3.**
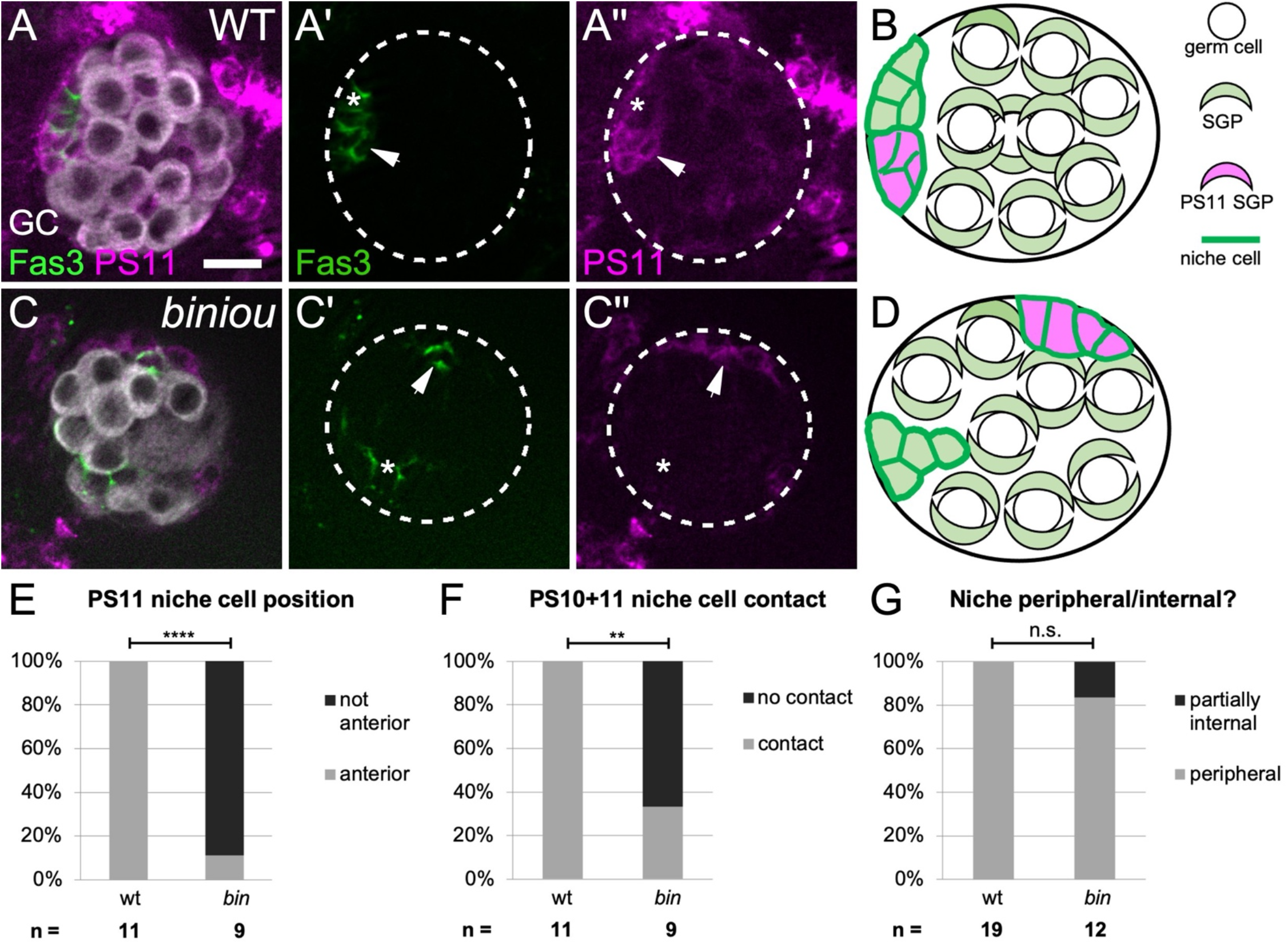
*biniou* is required for anterior movement of pro niche cells. (**A,C**) Stage 17 gonads expressing mcd8GFP in PS11 cells (magenta), and immunostained with Vasa (white, germ cells) and Fas3 (green, all niche cells). (**A**) A control, with a single anterior niche (left, green) containing cells deriving from both PS 10 (**A’**, green alone, asterisk) and PS 11 (**A’**, **A”**, magenta and green, arrow). (**C**) *biniou* mutant with dispersed niche cell aggregates (green). Anterior niche cells deriving from PS 10 (**C’**, green alone, asterisk) do not associate with PS 11-derived niche cells (**C”**, magenta and green, arrows). Ectopic PS11 niche cells were distinguishable from PS13 msSGPs, which do not express the niche cell marker Fas3. (**B,D**) Cartoons illustrating the distribution of PS11 niche cells in (**B**) control and (**D**) *biniou* mutants. (**E-G**) Quantifications comparing *biniou* mutants and sibling controls (Fisher’s exact test) by how often (**E**) PS 11-derived niche cells are located at anterior (p < 0.0001), (**F**) PS 11 niche cells contact anterior PS 10 niche cells (p < 0.0022), and (**G**) niche cells are located within 2 um of the gonad periphery. Scale bar, 10 um.

The second step of assembly requires anterior migration of pro-niche cells along the gonad periphery. A properly assembled niche is comprised of some cells that were initially specified near their final position since they derive from parasegment (PS) 10, and other cells that were specified more centrally and thus must migrate anteriorly as revealed by lineage tracing PS 11 cells (Anllo et al., 2019; DeFalco et al., 2008; Le Bras and Van Doren, 2006). After assembly, lineage-traced PS 11 niche cells labeled with mcd8GFP (magenta) and Fas3 (green), while PS 10 cells only labeled with Fas3 (green; **Fig 3A,** arrow**; Fig 3B**). When we lineage-traced PS 11 cells in *biniou* mutants most PS 11 niche cells remained in their original, more central positions, and were less frequently associated with PS 10 niche cells (**Fig 3C-F).** Since in the absence of Vm, PS 11-derived niche cells failed to reach the gonad anterior, this suggests a requirement for Vm signaling during the second step of niche assembly.

### Vm signaling results in *islet* expression in niche cells

Given the migratory steps in assembly, our prior finding of dispersed niches in gonads from *islet* (*tup*) mutants is revealing (Anllo et al., 2019) (**Fig 4A-C**), especially in light of its phenotypic similarity to *biniou* mutants, and combined *slit, pyr* and *ths* mutant (**Fig 1,2**). In fact, Islet protein was significantly enriched in niche cells (**Fig 4E; Fig S5, A-B**, and a minimal element from the *islet* enhancer region (Bataillé et al., 2020; Boukhatmi et al., 2014) was sufficient to drive GFP expression in niche cells (**Fig 4D**). Moreover, Islet protein accumulation depended on *biniou* (**Fig S5, C-D**), and on *slit*, or *pyr* and *ths* (**Fig 4F-I**). These results indicate that Slit and FGF signals, likely emanating from the Vm, act via islet to impact niche assembly.

**Figure 4.**
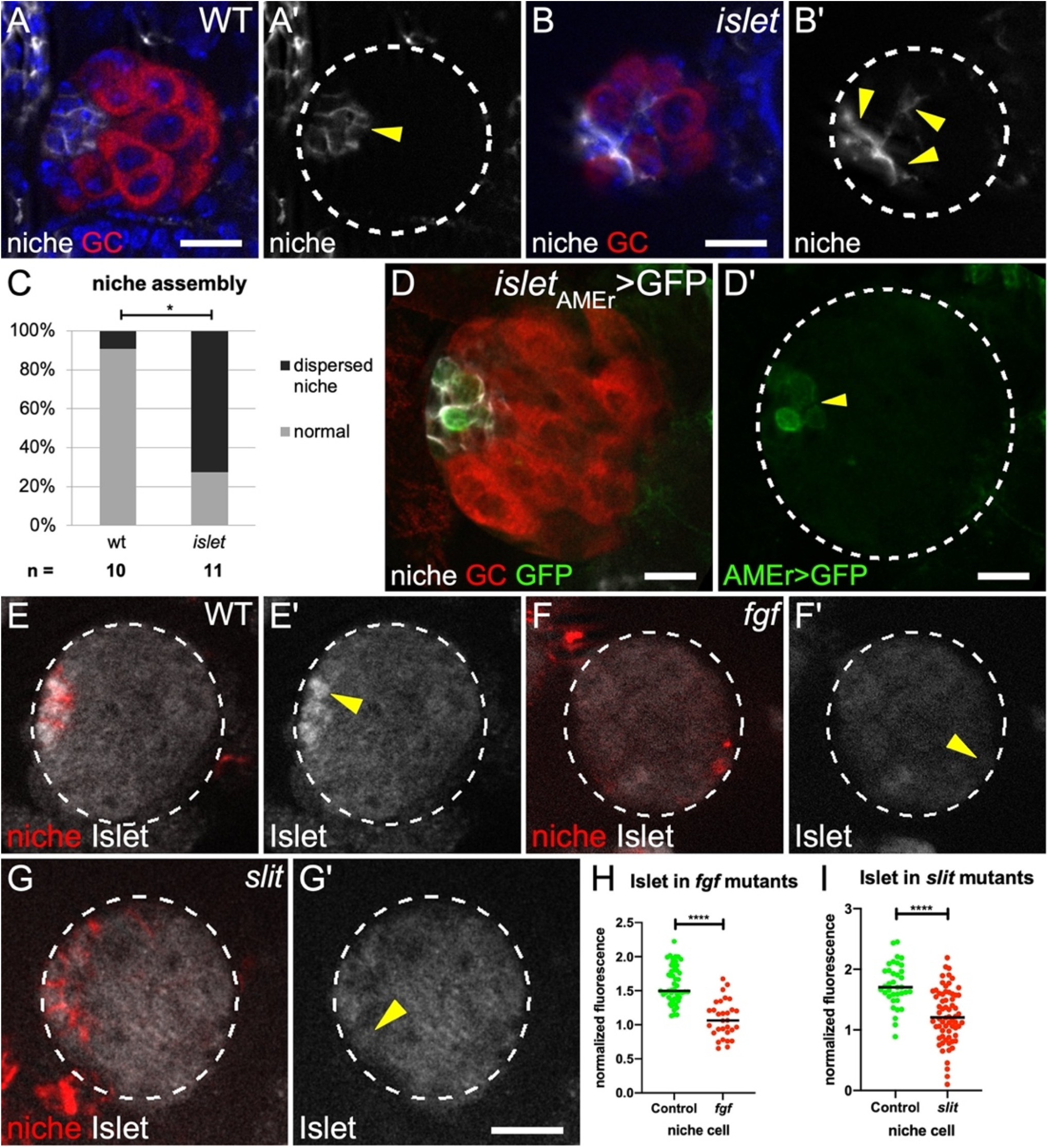
*islet* is expressed in niche cells in response to Vm signals. (**A**) Control and (**B**) *islet* mutant Stage 17 gonads immunostained with Vasa (red, germ cells), Fas3 (white, niche cells), and Hoechst (blue, nuclei). (**A’, B’**) Fas3 alone. (**C**) Quantification of niche assembly in *islet* versus sibling controls (p = 0.024, Mann Whitney test). (**D**) A gonad expressing GFP driven by the *islet* AMEr enhancer stained with Vasa (germ cells, red) and Fas3 (niche cells, white). (**D’**) *islet* AMEr driven GFP alone. (**E-G**) Stage 17 gonads immunostained for Islet (white), Fas3 (red, niche cells), and Vasa (not shown, germ cells). Gonad boundaries, dotted lines. Arrowheads, niche cells. (**E’-G’**) Islet alone. (**H-I**) Islet accumulation in niche cells from (**H**) *pyr* and *ths* removed *(fgf)* and (**I**) *slit* mutants, compared to sibling controls (p < 0.0001, Mann Whitney test). Scale bars, 10 um.

### Niche cells exhibit cytoskeletal polarity during assembly in response to Vm signaling

Since the cellular cytoskeleton is often polarized during migration, we examined the localization of F-actin during the later steps of niche assembly. Live-imaging revealed enrichment along niche-niche interfaces, as these cells began to associate with one another at the gonad anterior **(Fig 5A**, 0min). Interestingly, F-actin then re-polarized to niche-GSC boundaries as assembly completed (**Fig 5A**, 50min). Quantification in fixed tissue confirmed F-actin polarization during assembly (**Fig 5B**) and the shift afterwards (**Fig 5C**). Thus, niche cell cytoskeletal polarity is regulated during assembly.

**Figure 5.**
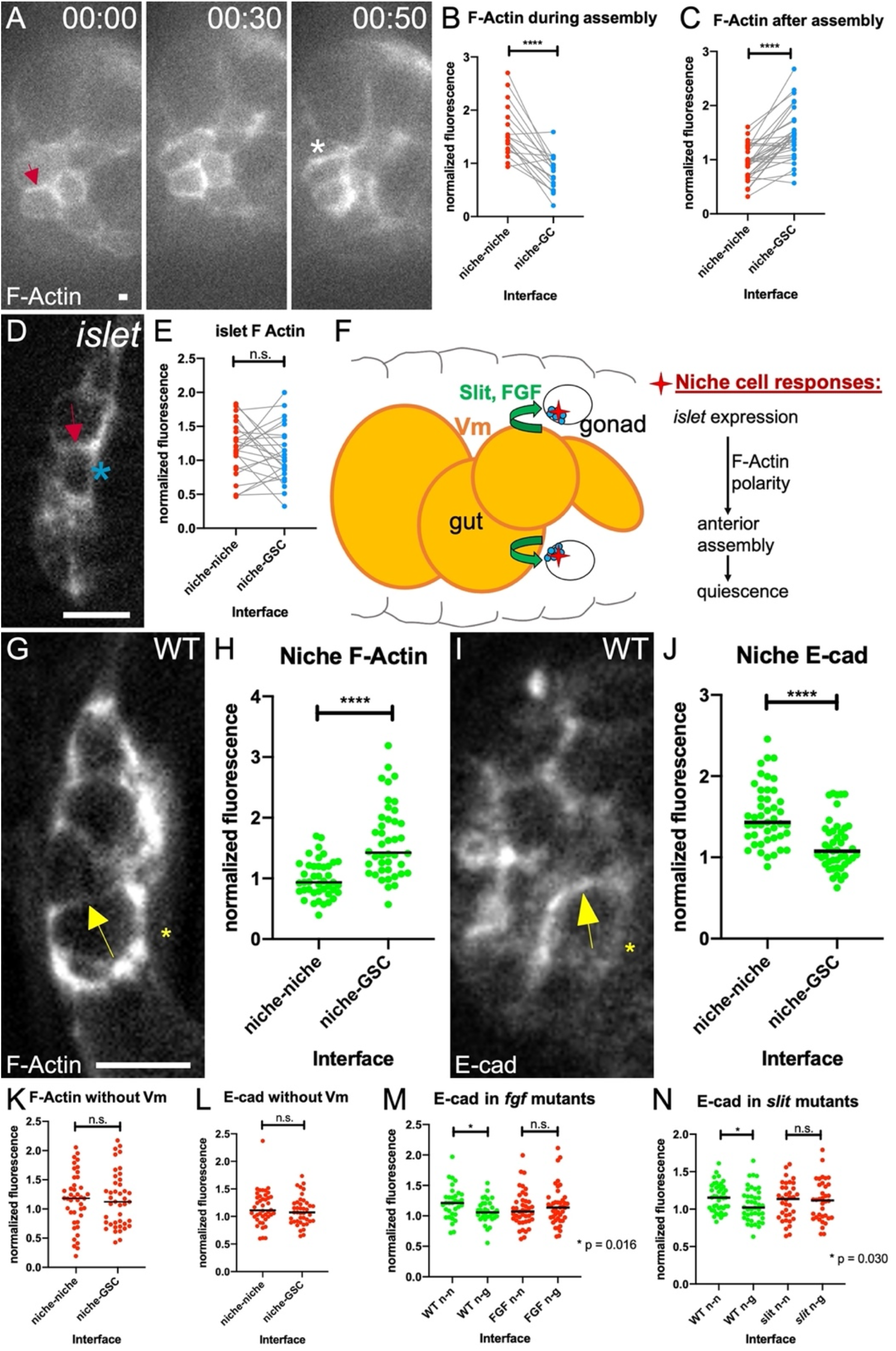
Niche cells are polarized during assembly. (**A**) Stills from a time-course of a gonad expressing *six4-* eGFP::moesin to label F actin in all SGPs. Factin accumulates at niche-niche interfaces when niche cells begin to associate (arrow), and later re-polarizes to niche-stem cell interfaces (asterisk). (**B,C**) Quantification of F actin accumulation at niche-niche or niche-germ cell interfaces (**B**) during and (**C**) after completion of niche assembly in fixed tissue (p < 0.0001, Wilcoxon test). (**D**) Niche cells in fixed tissue expressing a somatic cell F actin label, *six4-* eGFP::moesin in *islet* mutants in which niche cells have begun to associate, but have not completed assembly. (**E**) F actin accumulation at niche cell interfaces in *islet* mutants. (**F**) A model illustrating how Vm signals influence niche assembly. (**G,I**) Niche cells from Stage 17 control gonads (**G**) expressing *six4-* eGFP::moesin or (**I**) immunostained for E-cadherin. (**H**) F actin accumulation at niche-GSC interfaces versus niche-niche interfaces (p < 0.0001, Mann-Whitney test). (**J**) E-cadherin accumulates at niche-niche interfaces compared to niche-GSC interfaces (p < 0.0001, Mann-Whitney test). (**K-N**) Quantification of polarity loss in Stage 17 niches for (**K**) F-Actin in *biniou,* (**L**) E-cadherin in *biniou,* (**M**) E-cad with *pyr* and *ths* removed (*fgf*), or (**N**) E-cad in *slit* mutants (Mann Whitney tests). Asterisks, niche-GSC interfaces; arrows, niche-niche interfaces. Scale bar, 5 um.

Recognizing that *islet* encodes a transcription factor that regulates adhesion and guidance in the nervous system (Santiago and Bashaw, 2014; Santiago and Bashaw, 2017), we tested whether the cytoskeletal polarity observed for niche cells was disrupted in *islet* mutants. By quantifying polarity in mutants where niche cells had begun to associate with one another but had not completed assembly, we found that F-actin was not polarized (**Fig 5D,E**). Taken together, our results suggest that Vm signals Slit and FGF regulate *islet* expression in the niche, which impacts polarization of the F-actin cytoskeleton and also promotes assembly (**Fig 5F**).

Once formed it is known that the niche is enriched for cytoskeletal and adhesion proteins (Anllo et al., 2019; Le Bras and Van Doren, 2006). This prompted us to ask whether these components were polarized once assembly was completed. Indeed, both F-actin and E-cadherin were polarized, with F-actin enriched along niche-GSC interfaces compared to niche-niche interfaces, and E-cadherin enriched reciprocally, along nicheniche interfaces (**Fig 5G-J**). Interestingly, in Stage 17 gonads from *biniou* mutants neither F-actin nor E-cadherin were polarized in niche cells (**Fig 5K-L**). Similarly, gonads from *fgf* or *slit* mutants also failed to polarize E-cadherin (**Fig 5M-N**). These correlations suggest that without Vm signaling any associations among niche cells that do occur are not organized properly.

### Without Vm signaling, niche cells are functionally compromised and evade quiescence

We next tested whether niche assembly affected stem cell regulation. In the newly formed gonad, the niche recruits nearby germ cells to adopt stem cell fate, and orients stem cell divisions (Greenspan and Matunis, 2018; Hardy et al., 1979; Sheng et al., 2009; Sinden et al., 2012; Tanentzapf et al., 2007; Voog et al., 2008). One key niche-delivered signal, Upd, is known to activate the Stat pathway to higher levels among the first tier of germline cells adjacent to the niche (Anllo et al., 2019; Kiger et al., 2001; Leatherman and Dinardo, 2010; Leatherman and DiNardo, 2008; Sheng et al., 2009; Tulina and Matunis, 2001). We detected *upd* expression in niche cells in gonads from both control animals and *biniou* mutants (**Fig 1C,D**). As expected, in control gonads Stat protein was enriched in presumptive GSCs relative to neighboring germ cells (**Fig 6A-A’’, C**). In contrast, Stat enrichment was largely lost in gonads from *biniou* mutants, and from double mutants when *slit* and the *pyr* and *ths* fgf ligands were removed (**Fig 6B,B’, C**). These data suggest strongly that signaling from Vm affects niche function, and that a properly assembled niche might be required for robust signaling to the stem cells.

**Figure 6.**
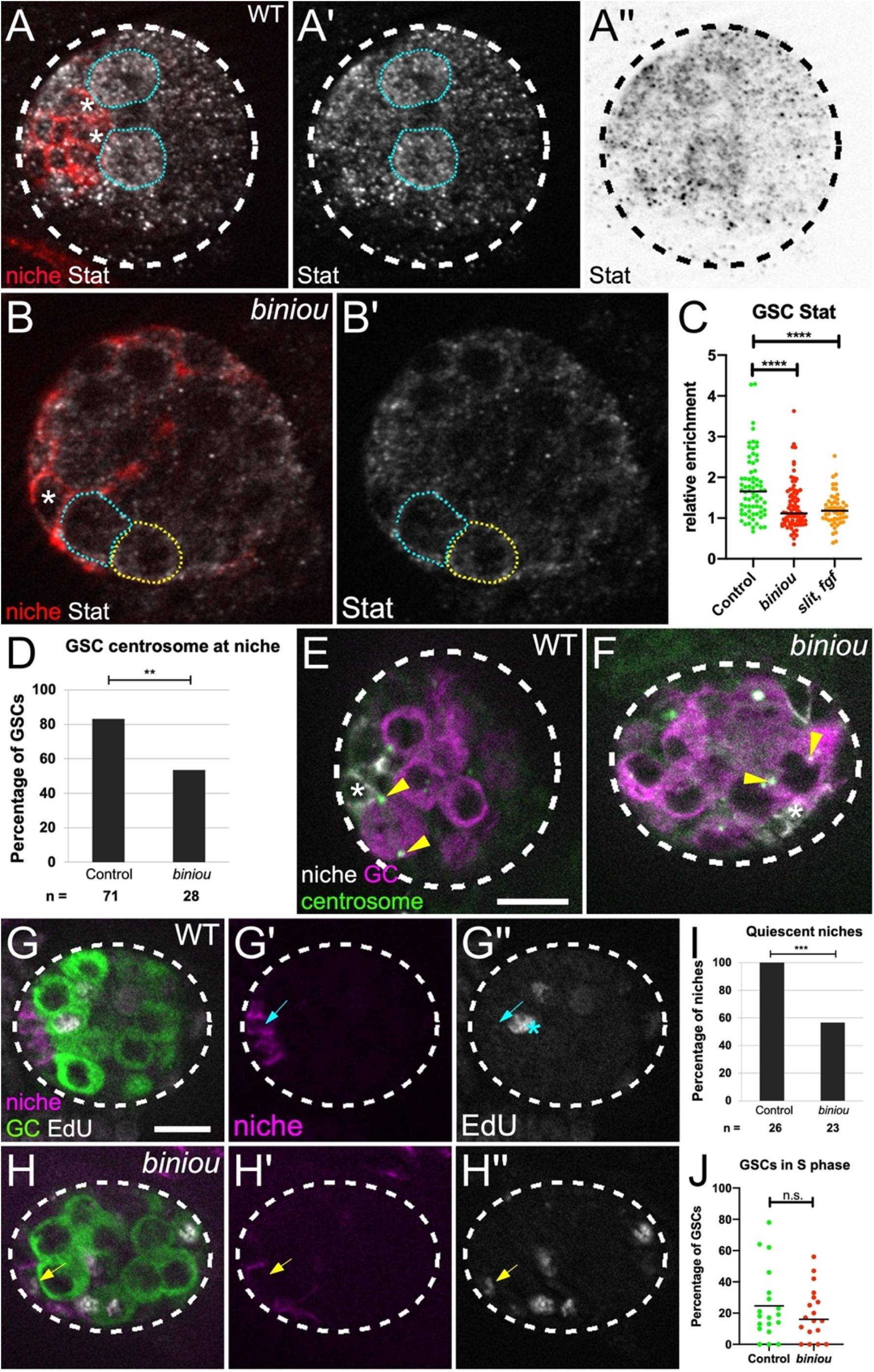
Niche assembly is required for niche function. (**A-B**) Stage 17 gonad, Stat (white), Fas3 (red, niche cells), and Vasa (germ cells; not shown). Niche cell, asterisk; GSCs, blue dotted line; neighboring germ cells, yellow dotted line. (**A’,B**’) Stat alone. (**A’’**) inverted Stat. (**C**) Stat accumulation in GSCs relative to neighboring germ cells in control, *biniou* or combined signaling mutant *(slit, pyr* and *ths*). (**D**) Quantification of control versus *biniou* mutants for percentage of GSCs with a centrosome at a GSC-niche interface (p = 0.004, Fisher’s exact test). (**E-F**) Stage 17 gonad, Gamma Tubulin (green, centrosomes), Fas3 (white, niche cells), and Vasa (magenta, germ cells). Arrowheads, GSC centrosomes; asterisk, adjacent niche cell. (**G-H**) Stage 17 gonads pulsed with EdU, fixed and immunostained. Merge shows EdU (white), Fas3 (magenta, niche), and Vasa (green, germ cell), along with single channel Fas3 (**G’,H’**), and EdU (**G’’,H’’**). **G’**) Control niche cell with no EdU incorporation (blue arrow), and **H’**) a *biniou* mutant niche cell incorporating EdU (yellow arrow). Asterisk, S phase GSC. (**I-J**) Quantification of control versus *biniou* mutants for: (**I**) Percentage of gonads with quiescent niches (p = 0.0001, Fisher’s exact test); **J**) S phase index of GSCs. Scale bars, 10 um.

Another key aspect of this niche is that it imposes oriented divisions on GSCs, such that daughter cells are displaced from the niche (Yamashita et al., 2003; Yamashita et al., 2007). In wild type testes, signals from the niche orient divisions perpendicular to the niche-GSC interface (Chen et al., 2018). To accomplish this orientation, one centrosome in the GSC remains near the interface with the niche, while the duplicated centrosome moves to the opposite pole of the GSC (Sheng et al., 2009; Yamashita et al., 2003) (**Fig 6E**). In contrast, in gonads from *biniou* mutants, both centrosomes in GSCs were often displaced from the interface with the nearby niche cell, suggesting a defect in centrosome anchoring (**Fig 6D,F**). These data again suggest functional defects in the niche in the absence of its proper assembly.

Strongly coupled to normal function of this niche is its quiescent state with respect to cell cycling (Greenspan and Matunis, 2018; Hetie et al., 2014). Indeed, EdU pulse-labeling showed that normal niche cells were quiescent (**Figure 6G,I**). In contrast, gonads with niche assembly defects exhibited many cycling niche cells (**Fig 6H,I**). This defect was selective in that the S-phase index for GSCs was similar to controls (**Fig 6J**). Thus, without proper assembly, pro niche cells fail to adopt their quiescent state.

Taken together, these results show defective niche signaling and behavior in the absence of Vm assembly cues, revealing that proper assembly is crucial to niche function.

## Discussion

We have shown that merely specifying niche cells is not sufficient for that niche to function. To adequately direct stem cell behavior, niche cells must be organized and positioned appropriately in the tissue. Prior live imaging suggested that niche placement was not simply congruent with the embryonic axes, but rather offset, tilted internally (Anllo et al., 2019). Here we reveal the visceral mesoderm (Vm) as the likely tissue required for that precision in niche placement, and we identify signals expressed in Vm that govern this process. We show that those signals are delivered early in gonadogenesis, and that in response, niche cells express the transcription factor *islet,* which plays a role in coordinating F-actin polarity in cells as they assemble into a niche that is functional and quiescent (see **Fig 5F**). Thus, this work identifies signaling, gene expression, and cell biological responses involved in regulating the assembly and proper positioning of the testis niche.

### Visceral mesoderm regulates development of the testis niche

We observed a striking dispersed niche phenotype in the absence of the transcription factor *biniou. biniou* is essential for the formation of Vm, and its expression had been reported as exclusive to the Vm (Zaffran et al., 2001). However, we observe Biniou accumulation in both the Vm and in gonadal SGPs (**Fig S2**). While this raised the possibility that Binou acted within SGPs, we identified two classes of ligands expressed in Vm, but not in SGPs that impact niche development in a manner similar to *biniou.* We also ruled out the possibility that the ligands control niche assembly by regulating *biniou* in SGPs because SGP accumulation of Biniou was unaffected in mutants where all ligands were removed (**Fig S2**). These data strongly suggest that the Vm directs anterior niche assembly.

The Vm and its signals appear to affect a specific step of niche assembly. We recently showed that pro-niche cells first extend protrusions to pull themselves out to the gonad periphery, then move anteriorly to associate and form an anterior cap on the gonad (Anllo et al., 2019). In *biniou* mutants, niche cells get to the periphery, but a subset of these cells cannot successfully arrive anteriorly (**Fig 3**). Niche cells derive from two separate clusters of mesodermal cells that only later associate as one niche (DeFalco et al., 2008; Le Bras and Van Doren, 2006). Pro-niche cells specified in PS 10 mesoderm are already located at what will be the gonad anterior, while those specified in PS 11 must migrate to reach the anterior. The lack of proper assembly without Vm signaling suggests either that pro niche cells cannot associate properly, or they cannot migrate. Our data suggests the latter. Without either *biniou* or the ligands Slit and FGF, we find that pro niche cells can still contact one another, likely as a result of sorting as niche cells upregulate adhesion proteins such as Fasciclin3, and E- and N-cadherin (Le Bras and Van Doren, 2006). However, our lineage tracing in *biniou* mutants revealed that PS 11-derived niche cells were almost never located at the gonad anterior (**Fig 3E**). We hypothesize that pro niche cells specified in different gonadal regions are not close enough to sort based on adhesion alone, and require Vm signals to enable movement to form a single niche. Note that Vm signals also act on PS10 niche cells as Islet is expressed in all niche cells and expression is lost in the absence of Vm signals.

### Vm signals are sent well before niche assembly

In many examples of cell migration, directive signals are active during the morphogenetic event (Montell, 2003; Scarpa and Mayor, 2016). We were surprised to find that the niche assembled normally in embryos that initially have Vm precursors but lose them prior to that time when the niche forms (**Fig 1**). From this data, we infer that Vm precursors emit the required signals early, significantly before niche assembly. One possibility is that early signaling induces a gene expression program in pro niche cells that enables an appropriate intrinsic cell response later in gonadogenesis. Indeed, we showed that before niche assembly, pro niche cells and Vm precursors directly intermingle and that *islet* gene expression is required downstream of Vm signaling. The identification of an *islet* cis-regulatory element sufficient for expression in the niche will help elucidate the circuitry involved in this induction event and establish whether it is a direct response to the signals defined here. Additionally, since *islet* itself is essential in niche assembly, its downstream targets will be of interest. In neurons, targets such as the DCC or Frazzled (Fra) receptor act in directing axons to their appropriate locations (Santiago and Bashaw, 2017). Perhaps such candidates might explain how *islet* induction contributes to niche assembly.

### Both Slit and FGF signals contribute to niche assembly and position

Our work has identified two signaling pathways important for niche assembly, suggesting that resiliency is built into the niche assembly process. While each pathway appears necessary for *islet* expression (Fig 4), which is important for niche assembly, the apparent dependence on signal dosage (**Fig S4**, **2**), and the fact that some niches can assemble in the absence of one pathway (see **Fig 2**) suggest that the pathways cooperate to ensure proper assembly and positioning. The contributions of Slit and FGF to niche assembly are reminiscent of the partially overlapping roles of EGFR and PVR in border cell migration during *Drosophila* oogenesis. The immediate downstream effectors of these pathways are unique, yet both EGFR and PVR converge on directing the migratory path of the border cells (Duchek et al., 2001).

In many cases, Slit and FGF function as directional guidance cues (Blockus and Chédotal, 2016; Friedl and Gilmour, 2009; Kadam et al., 2012). Changing the source of the cue in these instances alters the path of migrating cells (Jia et al., 2005; Sutherland et al., 1996). In niche assembly, we tested whether these pathways were acting in this manner. While we could not analyze niche assembly upon FGF misexpression, *slit* misexpression did yield gonads with properly assembled and positioned niches (**Fig 2K**). This result argues that Slit is in fact not acting as a directional cue during niche assembly. Slit protein is known to accumulate in nearby extracellular matrix in other systems (Isaacman-Beck et al., 2015; Xiao et al., 2011), and we likewise detected Slit accumulation near gonadal ECM, surrounding the periphery of the gonad (**Fig S3, G**). This apparently symmetric accumulation of Slit is consistent with our interpretation that Slit is not acting as a directional cue. Our data instead suggest that Slit provides for ‘competence,’ acting, for example, to enable pro niche cells to migrate, or licensing responses to as yet unidentified directional cues.

### Slit and FGF signaling affects cytoskeletal organization in pro niche cells

If Slit and FGF are eliciting responses prior to niche assembly, it is possible that they might regulate modulators of cell behavior required for assembly. Such regulators could be Islet targets. Cell movement often relies on asymmetric localization of cytoskeletal or adhesion proteins (Etienne-Manneville, 2008; Scarpa and Mayor, 2016; Vassilev et al., 2017). We showed that niche cytoskeletal polarity is normally well-organized during late stages of assembly, and that this organization is lost without *islet* (**Fig 5**). These data suggest that the niche assembly process depends on proper cytoskeletal polarization of pro niche cells in response to Vm signaling.

### Without proper assembly, niche cells function abnormally and evade quiescence

Niches are commonly found in a stereotypical position in each tissue. In mammals, the intestinal niche is within crypts, and the dermal papillae niche assembles at the base of the hair follicle (Wang et al., 2016). Such reproducibility in organization of niche cells suggests that proper niche assembly might be linked to its function, and our work reveals evidence of this link. We show that proper assembly of the testis niche is required to activate robust signaling in neighboring germ cells, and to orient stem cell divisions (**Fig 6A-F**), two crucial outputs of niche signaling. In the adult testis both of these outputs have been linked to intimate, cell biological organization at niche-stem cell interfaces (Chen et al., 2018; Inaba et al., 2015; Michel et al., 2011). Further, niche cells normally exhibit cell cycle quiescence (Greenspan and Matunis, 2018; Hardy et al., 1979; Voog et al., 2008), while promoting division of adjacent stem cells (Yamashita et al., 2003). We reveal an association between initial niche assembly and quiescence. It is clear that quiescence is important to the biology of the testis, as aberrantly dividing niche cells can generate extraneous niches located away from the testis tip and even lead to niche decay (Greenspan and Matunis, 2018; Herrera et al., 2021). There is evidence for feedback from other adult somatic cells in maintaining niche quiescence, but how niche cells first enter quiescence is unknown. All embryonic SGPs are cycling prior to niche formation, but fully assembled niche cells withdraw from cycling (**Fig 6G, I**), and assembly appears correlated with withdrawal (**Fig 6H,I**). Whether and how niche-niche cell contact and the tight regulation of cytoskeletal organization during assembly, including the asymmetric enrichment of E-Cadherin in assembled niches, is related to quiescence will require further study.

Together, our work identifies extrinsic signaling, and intrinsic gene expression and cell biological responses involved in governing niche cellular organization and position, which are integral to proper function of the testis niche.

## Supporting information

Reagent Table

Genotype Table

## Acknowledgements

We thank the Bloomington Drosophila Stock Center (NIH P40OD018537) and R. Lehmann, E. Bach, A. Holz, and M. Frasch for antibodies and stocks. Thanks to G. Bashaw, M. Granato, K. Nelson, G. Vida, B. Warder and especially K. Lenhart for comments.

## Author contributions

Conceptualization, L.A. and S.D.; Methodology, L.A. and S.D.; Validation, L.A.; Formal Analysis, L.A.; Investigation, L.A.; Resources, S.D.; Data Curation, L.A. and S.D.; Writing – Original Draft, L.A.; Writing – Review & Editing, L.A. and S.D.; Visualization, L.A.; Supervision, S.D.; Project Administration, L.A. and S.D.; Funding Acquisition, L.A. and S.D.

---

**Figure S1.**
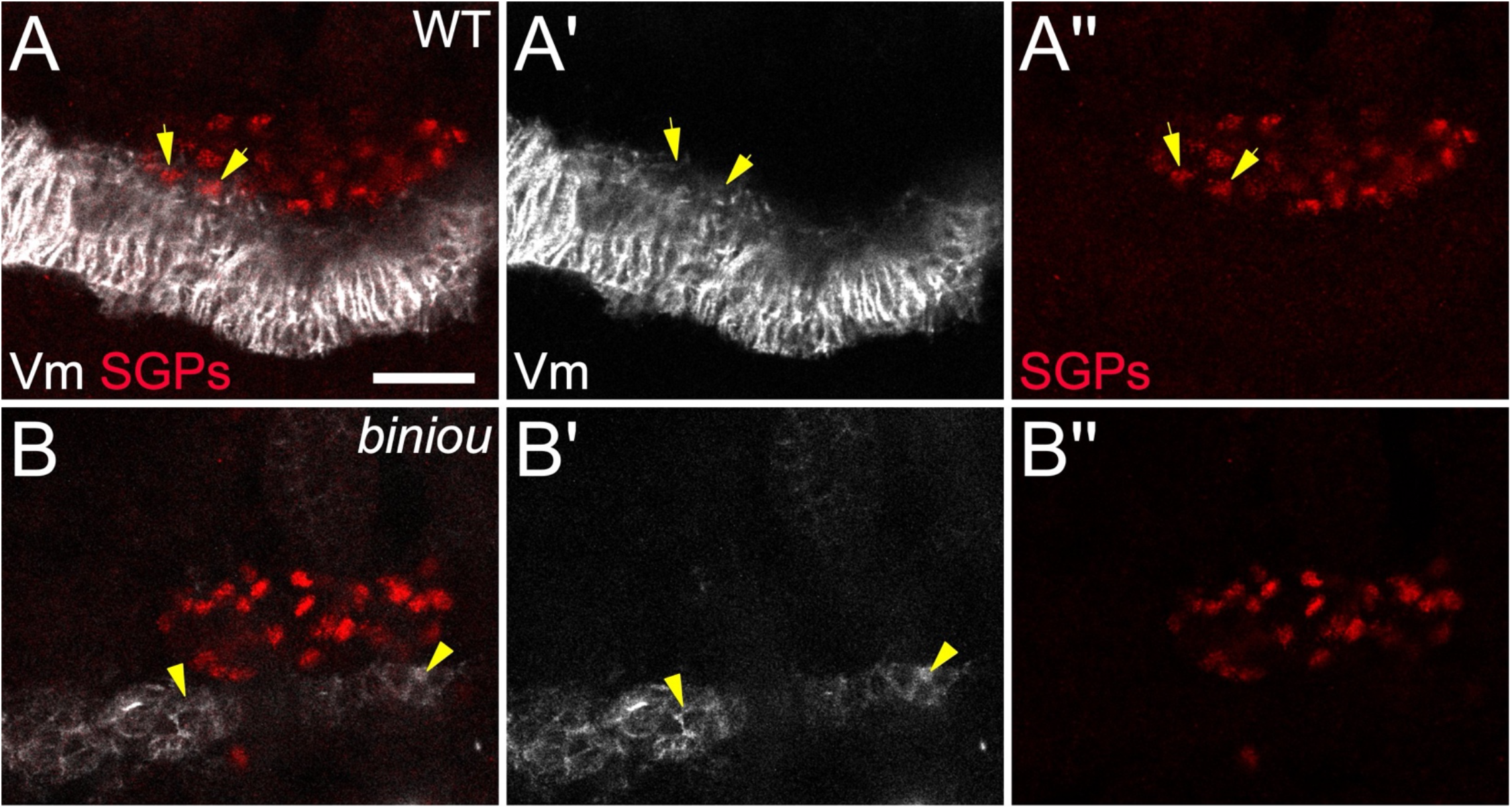
*biniou* mutant SGPs do not contact Vm precursors. SGPs (Traffic jam, red) that are still coalescing, and neighboring Vm (Fas3, white) in (**A**) control and (**B**) *biniou[R22]* mutant embryos. (**A**) Arrows indicate two SGP nuclei that are directly adjacent to Fas3 positive Vm tissue. In (**B**), there is less Fas3 positive tissue, expected for *biniou* mutants. Vm tissue is near the gonad, but not contacting SGP nuclei (arrowheads). Scale bar, 20 um.

**Figure S2.**
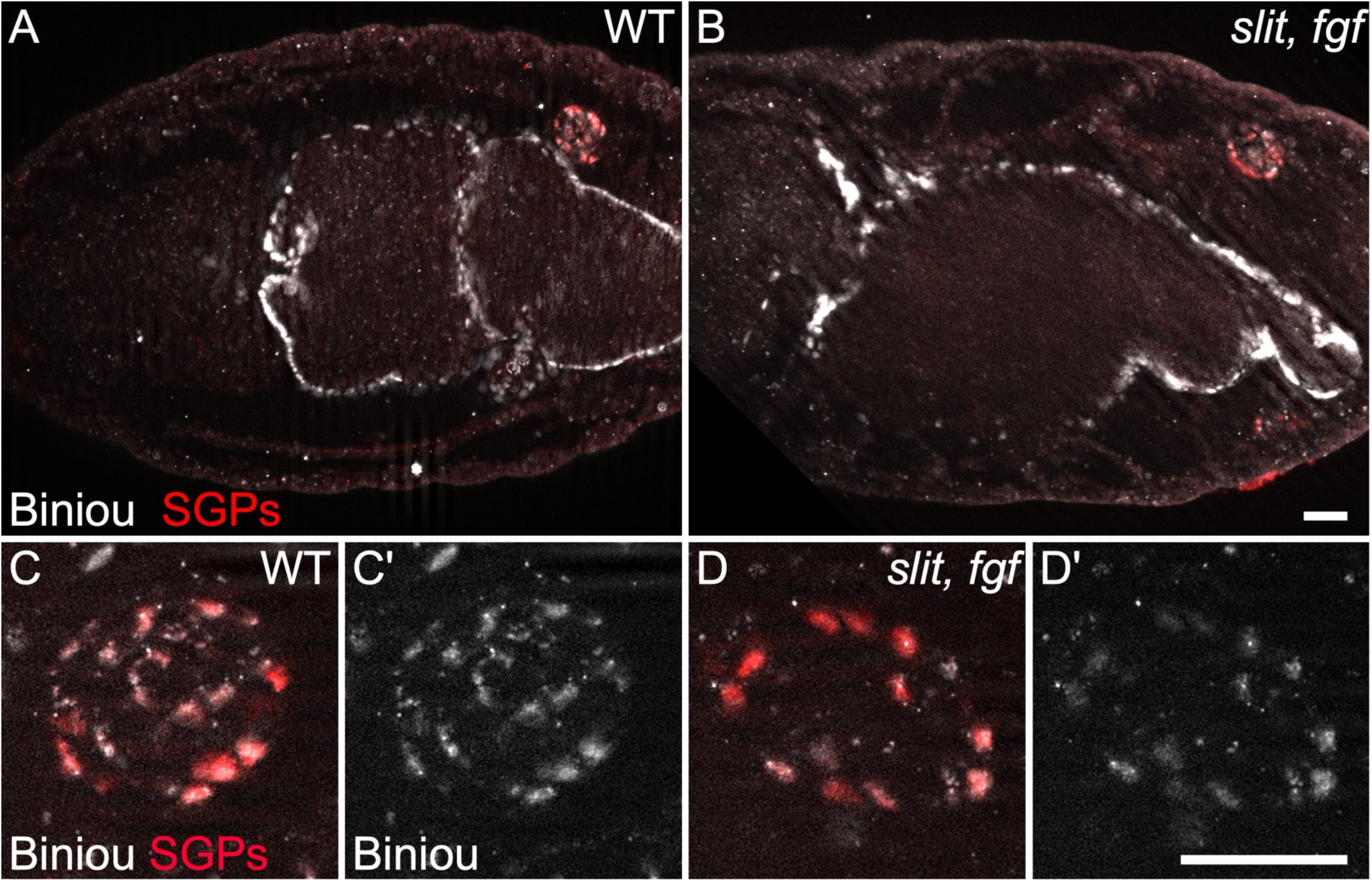
SGPs express *biniou*. Stage 16 (**A-B**) embryos, and (**C-D**) gonads immunostained for Biniou (white) and Traffic jam (red, SGPs). SGPs from (**A,C**) wild type or (**B,D**) combined mutants with *slit,* and *pyr* and *ths* removed *(fgf)* express *biniou.* Scale bars, 20 um.

**Figure S3.**
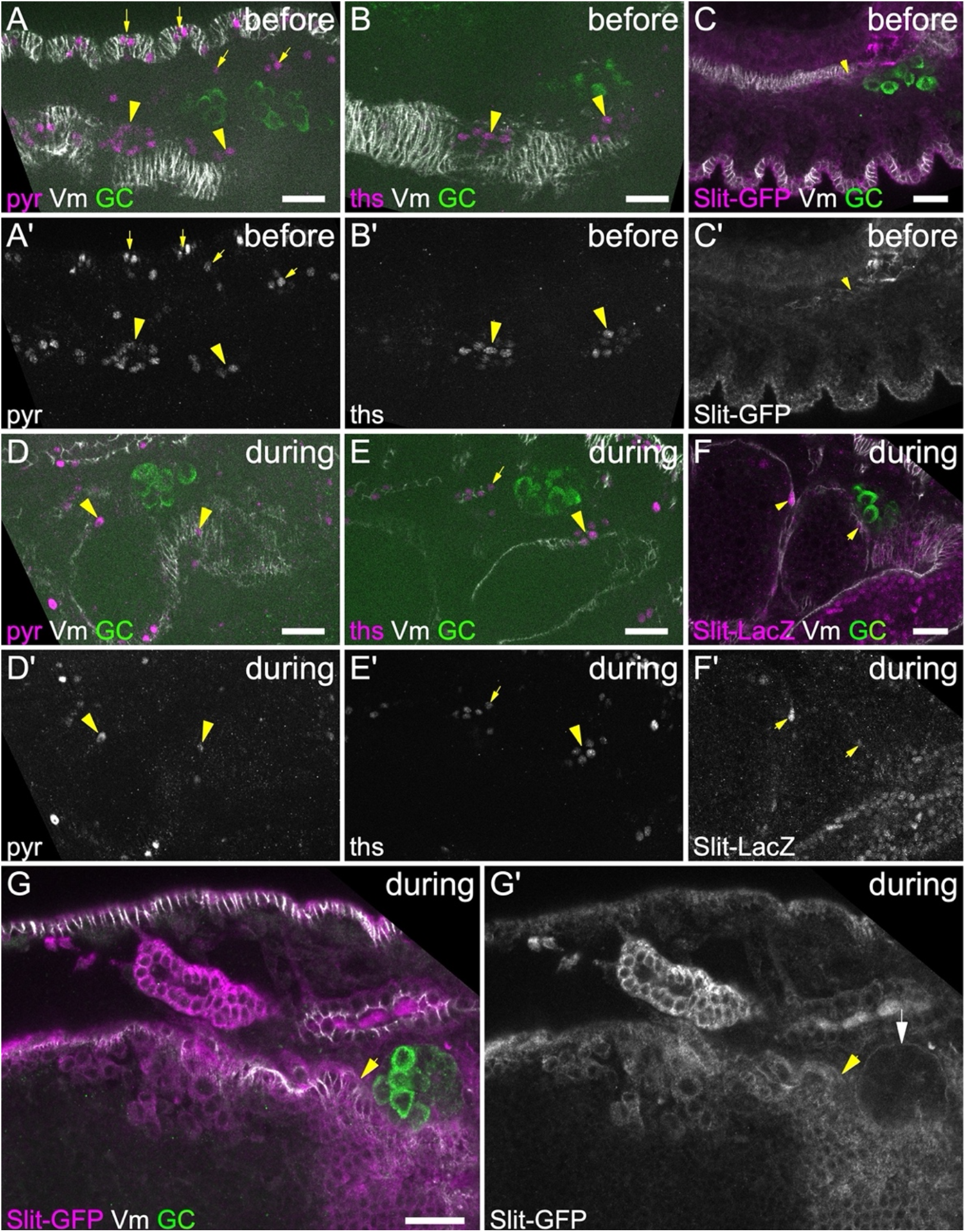
FGF and Slit are expressed in visceral mesoderm. (**A-G**) Embryos immunostained for Vasa (green, germ cells), and Fas3 (white) to show visceral mesoderm and other tissues, including the overlying epidermis, and the tubular hindgut and foregut. (**A,D**) Embryos express *pyr*-Gal4 driving UAS-red Stinger (magenta) to show FGF ligand expression (**A**) before or (**D**) during niche assembly. (**A, A’**) Arrowheads show *pyr*-expressing cells within Vm. Arrows reveal a metameric pattern of Pyr in overlying epidermis and some underlying mesoderm. At this stage SGPs are intermingled with germ cells where little or no Pyr is detectable. (**D, D’**) Pyr is clearly detectable among Vm cells (arrowheads) and the hindgut, but not among germ cells. (**B,E)** Embryos express *ths*-Gal4 driving UAS-red Stinger (magenta) to show expression of the second, redundant FGF ligand. As with *pyr,* we observe expression in the Vm (arrowheads) and some neighboring mesodermal cells (arrows), but not among germ cells. (**C,F**) Embryo expressing Slit::GFP or Slit-LacZ (magenta) (**C**) before or (**F**) during niche assembly. (**C, C**’) Arrowheads indicate clusters of Vm cells expressing Slit::GFP including those near germ cells. Slit expression is also observed in cells neighboring the gonad that do not derive from visceral mesoderm. (**F, F’**) Occasional Slit-expressing Vm cells are still present (arrow), but little or no expression is detectable among germ cells. (**G, G’**) Slit-GFP protein accumulates near gonadal extra-cellular matrix (ECM, white arrow) during niche assembly. Nearby Vm cells expressing Slit are indicated with a yellow arrow. Scale bars, 20 um.

**Figure S4.**
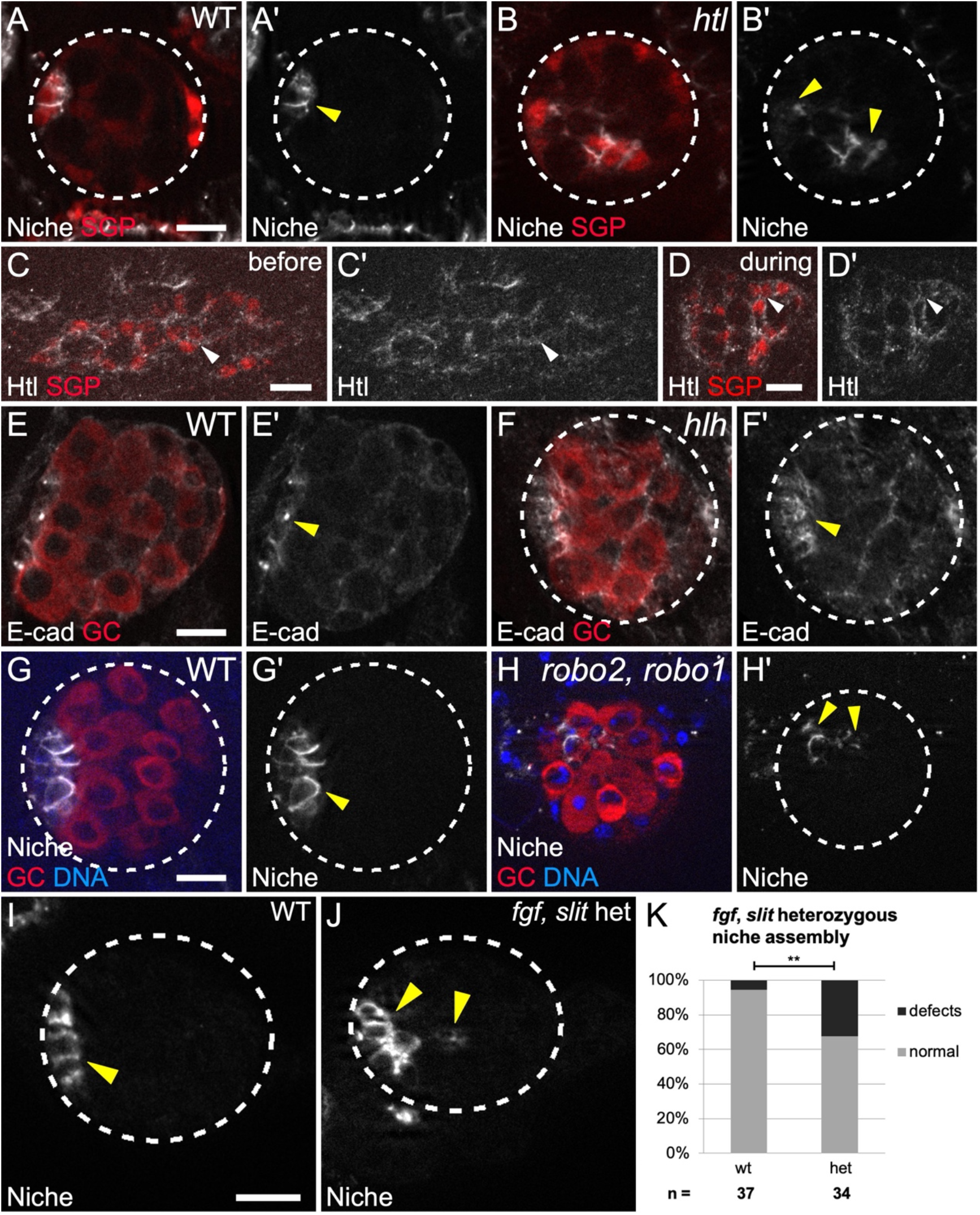
FGF and Slit receptors and dosage are important for niche assembly. (**A-B**) Stage 17 gonads expressing *six4*-nls-eGFP (SGPs, red) immunostained for Fas3 (niche cells, white). (**A’,B’**) Fas3 alone. (**A**) Wild type gonads have a single anterior niche, while (**B**) *htl* receptor mutants exhibit dispersed niche cell aggregates. (**C,D**) Htl-mCherry expressing gonads (**C**) before and (**D**) during assembly immunostained for RFP (Htl, white) and TJ antibody (SGPs, red). Neighboring SGPs show Htl localization at cell boundaries. (**E,F**) Stage 17 gonads stained with Vasa (red, germ cells) and E-cadherin (white, enriched in the niche). (**E’,F’**) Ecad alone. Both (**E**) controls and (**F**) *hlh54f* cvm mutants have an anterior niche. (**G,H**) Stage 17 gonads immunostained for Vasa (germ cells, red), Fas3 (niche cells, white), and Hoechst (DNA, blue). (**G’,H’**) Fas3 alone. (**I-J**) Stage 17 gonads stained for Vasa (germ cells, not shown), and Fas3 (niche cells, white). (**I**) Controls have a single anterior niche, in contrast to (**J**) *slit, FGF* heterozygotes which show dispersed niche cells. **(K)** Quantification of niche assembly in *slit, fgf* heterozygotes (p = 0.005, Fisher’s exact test). In all panels: yellow arrowheads, niche cells; dotted lines, gonad boundaries; scale bars, 10 um.

**Figure S5.**
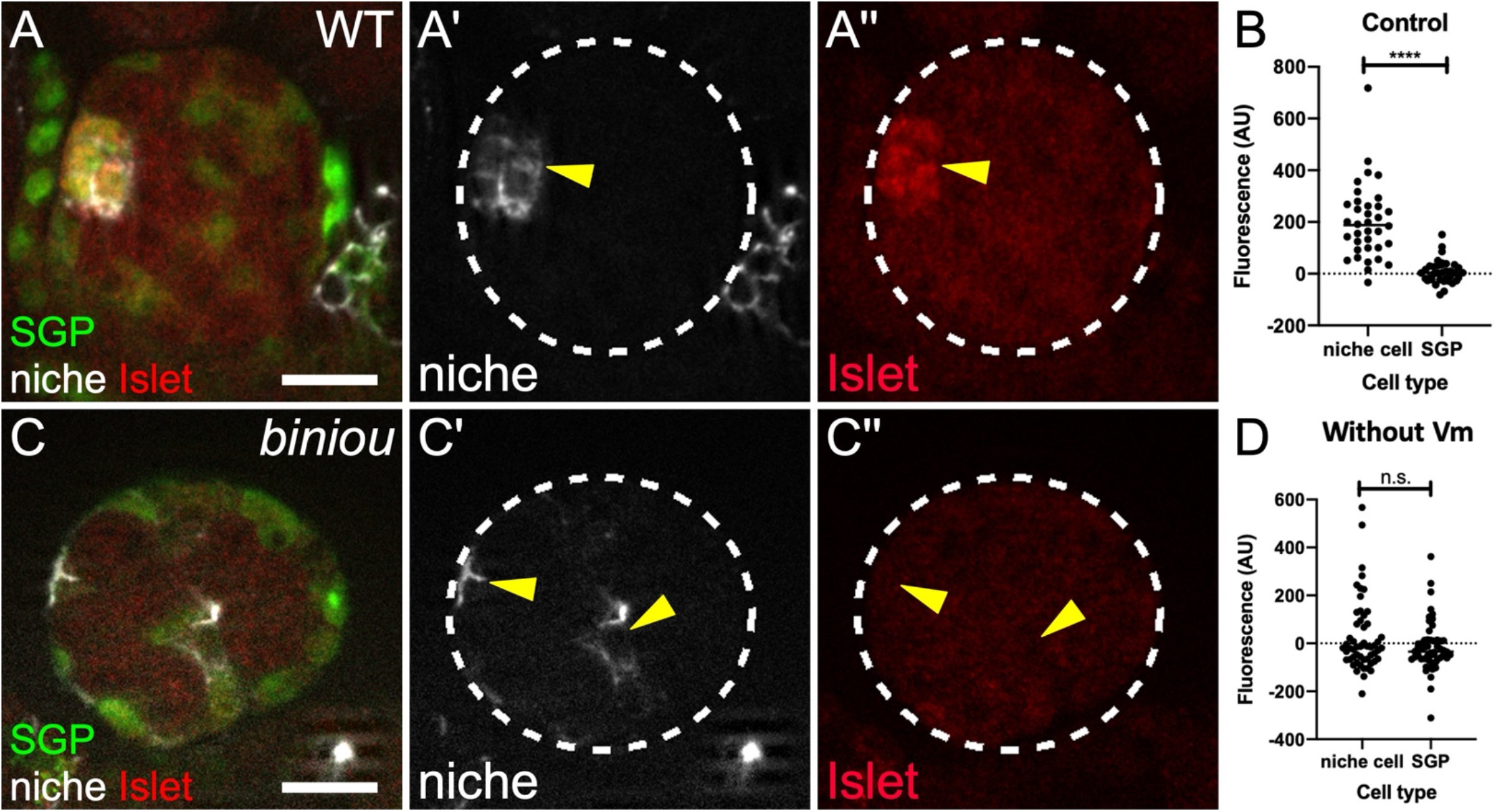
*islet* expression depends on *biniou*. (**A,C**) Stage 17 *six4nlsGFP* gonads immunostained for GFP (green, SGPs and other mesoderm), Fas3 (white, niche cells), and Islet (red). (**A’,C’**) Fas3 alone. (**A’’,C’’**) Islet alone. (**A**) Control gonads express Islet in niche cells, while (**C**) *biniou* mutants do not. (**B,D**) Islet accumulation in Fas3 positive niche cells, compared to non-niche SGPs in (**B**) control (p < 0.0001, Mann Whitney test) and (**D**) *biniou* mutants.

## Methods

### RESOURCE AVAILABILITY

#### Lead contact

- Further information and requests for resources and reagents should be directed to and will be fulfilled by the lead contact, Stephen DiNardo.

#### Materials availability

- This study did not generate new unique reagents.

#### Data and code availability

- All data reported in this paper will be shared by the lead contact upon request.
- This paper does not report original code.
- Any additional information required to reanalyze the data reported in this paper is available from the lead contact upon request.

### EXPERIMENTAL MODEL AND SUBJECT DETAILS

#### *Drosophila* Stocks

All *Drosophila* lines used are listed in the Key Resources Table (KRT). *slit[2]* is a null allele with no detectable protein product (Battye et al., 2001; Nusslein-Volhard et al., 1984). To remove *pyr* and *ths* together, we used a small chromosomal deficiency, Df(2R)BSC25, which completely deletes the genes encoding both ligands (Stathopoulos et al., 2004). *jeb[weli]* mutants lack visceral muscle founder cells (Stute et al., 2004), and the small chromosomal deficiency, Df(2R)BSC699 uncovers *jeb. hlh54f[delta598]* mutants lack caudal visceral mesoderm (Ismat et al., 2010). *htl[AB42]* is a null allele of the FGF receptor *heartless* (Gisselbrecht et al., 1996). *robo2[X123]* and *robo1[GA285]* are null alleles of the Slit receptors *robo2* and *robo1* (Evans et al., 2015; Kidd et al., 1998).

#### Sex Identification and Genotyping

Gonad sex identification was accomplished as described by Anllo and colleagues (Anllo et al., 2019). We used Vasa antibody staining to identify larger male gonads, and male specific SGPs (msSGPs). Vasa antibody labels both germ cells and msSGPs, and we identified msSGPs using Vasa antibody alongside a DNA stain to indicate small, Vasa positive nuclei. Sibling controls were distinguished from homozygous mutants by using fluorescent balancer chromosomes (TM3, P{w[+mC]=Gal4-twi.G}2.3, P{UAS-2xEGFP} AH2.3, Sb[1], Ser[1], FBst0006663; TM6, P{Dfd-EYFP}, Sb, Hu, e; or CyO, P{Dfd-EYFP}).

### METHOD DETAILS

#### Embryonic gonad dissection and Immunostaining

Dissections and immunostaining were performed as previously described (Anllo et al., 2019). Embryos were collected and aged 22-25 hours in a humidified chamber at 25 degrees C for late stage 17 embryos. For younger embryos still undergoing late stages of niche assembly, embryos were aged 22.5-24.5 hours at 23 degrees C. Primary antibodies were used overnight at 4 degrees C. Secondary antibodies were used at 3.75 ug/mL (Alexa488, Cy3, or Alexa647; Molecular Probes; Jackson ImmunoResearch) for 1-2 hr at room temperature. DNA was stained with Hoechst 33342 (Sigma) at 0.2 ug/mL for 5 min.

We used rabbit antibody against Vasa 1:5000 (gift from R. Lehmann, NYU), and RFP 1:500 (Abcam, ab62341); goat antibody against Vasa 1:200 (Santa Cruz, dC-13, now discontinued); mouse antibody against Fasciclin III 1:50 (DSHB, 7G10), Islet 1.5:100 (DSHB 40.3A4; *Drosophila* Tailup), and Gamma Tubulin 1:200 (Sigma, GTU-88); rat antibody against DE-cadherin 1:20 (DSHB, DCAD2); guinea pig antibody against Traffic jam 1:10,000 (gift from D. Godt); chick antibody against GFP 1:1000 (Aves Labs, GFP-1020); and rabbit antibody against Biniou 1:100 (gift from E. Furlong). Images of fixed samples were acquired on a Zeiss Imager with Apotome using a 40x, 1.2 N.A. lens or a 20x, 0.8 N.A. lens; or on a Zeiss LSM 880 Confocal with Airyscan and Fast Airyscan, 40x, 1.2 N.A. lens.

#### Identification of niche position

To confirm the position of niche cells relative to the anterior-posterior axis of the gonad, we used the position of the male specific somatic gonadal precursor cells (msSGPs) to denote the gonad posterior (DeFalco et al., 2003). msSGPs are visible to a trained eye in many stains, and can be detected as a cluster of cells distinct from the germ cells at the posterior pole of the gonad with Vasa immunostaining (Anllo et al., 2019; Sheng et al., 2009). Because the embryonic gonad has a spherical shape, we confirmed an anteriorly positioned niche by its location at the pole of the gonad roughly 180 degrees opposite to where the msSGP cluster resides.

#### Niche phenotypic characterization

Normal niches were located in a single grouping of cells at the gonad anterior, with a smoothened boundary. Dispersed niches included a range of phenotypes, including cases where multiple distinct niche cell groupings were present within a single gonad, and cases where a single niche cell grouping formed with highly irregular boundaries.

#### *in vivo* live imaging

Live imaging was performed as previously described (Anllo et al., 2019; Ong et al., 2019). Images were acquired with a Leica DM16000 B spinning disk confocal with a 63 x 1.2 N.A. water immersion objective, using an EMCCD camera (Andor iXon 3 897E or Hamamatsu photonics, model C9100-13) controlled by Metamorph software. Z stacks were taken at 5-minute intervals, with 36 1 um z-slices through the gonad.

#### Slit and FGF ligand overexpression

The *twi-Gal4* driver was use to over-express either *UAS-sli.D* or *UAS-ths289.22* in all mesodermal cells. Embryos were collected for 2-3 hours at 29 degrees C, and were aged 15-18 hours at 29 degrees prior to dissection. Just prior to dissection, UAS-eGFP expression was used to distinguish and sort embryos that carried the *twi*-Gal4 driver from sibling controls.

#### EdU Pulse experiments

EdU pulse experiments were performed using the Click-iT EdU Plus kit (Molecular Probes, c10640) (Salic and Mitchison, 2008). Immediately after dissection, tissue was incubated in 10 uM EdU in *Drosophila* Ringers solution for 30 minutes at room temperature. Tissue was then fixed for 15 min in 4% PFA at room temperature.

The azide reaction to couple EdU to alexa647 was performed either prior to, or after antibody staining. Copper catalyst was used at a concentration of 4 nM.

### QUANTIFICATION AND STATISTICAL ANALYSIS

#### Counting niche cells

Niche cells were identified using the niche-cell specific Fasciclin III immunostain. Niche cell nuclei were counted, using either Hoechst DNA stain, or Traffic Jam nuclear stain, as a marker. The ImageJ Cell Counter plugin was used to record counted niche cells. A Mann-Whitney test was used to determine significance of p < 0.05.

#### Quantification of *islet* expression in *slit,* and *pyr* and *ths* removed *(fgf)* mutants

To quantify *islet* expression in Vm ligand mutants, we stained gonads with Islet antibody and used ImageJ to measure the mean gray value fluorescence intensity within regions of interest (ROIs). We selected ROIs including a circular region within somatic cell nuclear boundaries, using Hoechst stain as a marker. For each gonad, 3 niche cell ROIs were measured for Islet expression. An ROI devoid of tissue was selected in a region adjacent to the gonad to determine background fluorescence. Background fluorescence was subtracted from measured niche cell values. Each background-subtracted value was normalized to the mean Islet fluorescence for the gonad, measured at a Z slice including the niche. Mann-Whitney tests were used to determine significance of p < 0.05.

#### Quantification of *islet* expression in *biniou* mutants

To quantify *islet* expression in *biniou* mutants, we stained gonads with Islet antibody and used ImageJ to measure the mean gray value fluorescence intensity within regions of interest (ROIs). We selected ROIs including a circular region within somatic cell nuclear boundaries, using *six4*nlsGFP as a marker. ROIs were in a single Z plane in which the relevant nucleus was in focus. For each gonad, 3 niche cells and 3 non-niche SGP ROIs were measured for Cy3 Islet and for GFP nuclear marker fluorescence. An ROI selected to encompass the unlabeled region of a single germ cell within each gonad was used to determine background fluorescence. Background fluorescence was subtracted from measured values.

To control for possible bleed-through of GFP nuclear marker into the Cy3 Islet channel, we first measured the amount of Cy3 signal that could be accounted for by GFP bleed-through. We plotted the ratio of Cy3 to GFP fluorescence intensity in gonads that were not stained with Islet antibody, and thus should not have any Islet Cy3 signal. This plot determined that Cy3 signal resulting from bleed-through averaged 7% of the GFP signal intensity for each ROI. Thus, in addition to background subtraction, we also subtracted 7% of the GFP signal values from Cy3 values to obtain our final measurements of Cy3 Islet signal. These values were plotted. Mann-Whitney tests were used to determine significance of p < 0.05.

#### Quantification of normalized F actin and E-cadherin fluorescence

To visualize F actin we imaged gonads expressing a GFP-labeled F actin binding protein in the somatic cells, *six4*-eGFP::moesin (Sano et al., 2012). E-cadherin was visualized by immunostaining with an antibody against E-cadherin (DSHB). For all experiments, gonads were dissected and immunostained either with an antibody against GFP, or E-cadherin. Niche interfaces were identified with a Fas3 immunostain. F actin or E-cadherin fluorescence intensity at niche-niche and niche-GSC interfaces was quantified using ImageJ to trace interfaces, and report mean gray values. Background fluorescence was measured as the mean gray value of a line traced where no tissue was present for E-cadherin experiments, or within a germ cell for F actin, as germ cells do not express *six4*-eGFP::moesin. After background subtraction, fluorescence intensity was normalized by taking the ratio of each interface measurement to the average of all interfaces within that gonad. Normalized values were then plotted, and data was analyzed using a Mann-Whitney test.

#### Centrosome position quantification

Centrosome position was visualized with immunofluorescence against Gamma tubulin to label pericentriolar material. GSCs were scored for centrosome position if they had already undergone centrosome duplication. We quantified how often one of the two centrosomes was located closer to the adjacent niche than to other neighboring cells. Those GSCs with a centrosome located near the niche-GSC interface were scored as appropriately positioned. GSCs in Vm mutants often failed to maintain a centrosome near the niche. GSCs in Vm mutants that made contact with niche cells at multiple points around their periphery were scored as normal if a centrosome was close to one of these niche-GSC contacts. Data was analyzed using Fisher’s exact test.

#### Quantification of Stat accumulation

To quantify Stat accumulation, we stained gonads with Stat antibody (E. Bach, 1:1000) and used ImageJ to measure the mean gray value fluorescence intensity within regions of interest (ROIs). We selected ROIs including a circular region to sample germ cells, using Vasa immunofluorescence as a marker to delineate cell boundaries. For each gonad, we sampled 5 GSCs and 3 neighboring germ cells. After background subtraction, we measured the ratio of Stat accumulation within each GSC relative to the neighboring germ cell average for that gonad. Relative Stat enrichment values were plotted for each GSC. We obtained measurements on sibling controls and in mutants for *biniou,* or for combined *slit* and *fgf*-removed mutants. Mann-Whitney tests were used to evaluate comparisons.

